# GpABC: a Julia package for approximate Bayesian computation with Gaussian process emulation

**DOI:** 10.1101/769299

**Authors:** Evgeny Tankhilevich, Jonathan Ish-Horowicz, Tara Hameed, Elisabeth Roesch, Istvan Kleijn, Michael PH Stumpf, Fei He

## Abstract

Approximate Bayesian computation (ABC) is an important framework within which to infer the structure and parameters of a systems biology model. It is especially suitable for biological systems with stochastic and nonlinear dynamics, for which the likelihood functions are intractable. However, the associated computational cost often limits ABC to models that are relatively quick to simulate in practice. We here present a Julia package, GpABC, that implements parameter inference and model selection for deterministic or stochastic models using i) standard rejection ABC or ABC-SMC, or ii) ABC with Gaussian process emulation. The latter significantly reduces the computational cost.

URL: https://github.com/tanhevg/GpABC.jl

## Introduction

Parameter estimation and model selection are the central tasks in reverse engineering of cellular systems. Although identifying the best parameter value or model structure is of obvious interest, so too is the evaluation of the estimation uncertainty, which has contributed to the rising popularity of Bayesian inference. Much of the work has focused on Approximate Bayesian Computation (ABC) for both parameter and model inference^1, 2^, since the likelihood of nonlinear biological models is often intractable. A number of improvements on the basic ABC rejection scheme have been proposed, employing either Markov Chain Monte Carlo (ABC-MCMC)^3^ or sequential Monte Carlo (ABC-SMC)^1, 4^. However, even with these optimisations the number of time-consuming model simulations required can still easily reach the millions.

A further speed-up of ABC can be achieved by employing emulation techniques, where a mapping between the parameters and the approximated likelihood of a complex model (i.e. discrepancy between the model outputs and measurement data) is built using statistical regression models. Only a small number of simulations is required to train the emulator, which can then be used to predict the model outputs (or the discrepancy) for other parameters with a significantly lower computational cost. Accelerating ABC or MCMC with emulation has been proposed using either local regression or Gaussian process (GP) regression^5–7^. The GP-based approach has gained more traction recently, partly due to its inherent ability to quantify uncertainty and partly due to increased availability of computational resources.

A number of ABC packages have been published in the literature (for a review see^8^), however an ABC package with a focus on emulation is still lacking. Here, we present an extensible Julia package, GpABC, which implements rejection ABC and ABC-SMC with GP emulation for parameter and model inference in deterministic and stochastic models. Below we introduce the basic concepts of ABC and GP regression, and provide an overview of the package. Details of the algorithms, user guide, documentation and examples are available online at https://github.com/tanhevg/GpABC.jl.

## Materials and methods

### Approximate Bayesian computation (ABC)

The simplest version of ABC, ABC rejection, proceeds as follows: (i) sample a parameter vector, or particle, *θ* from the prior distribution; (ii) simulate a dataset *D* from the model given *θ* and compute summary statistics *S*(*D*|*θ*) if the dataset is high-dimensional; (iii) compute a distance *d* that quantifies the discrepancy between *S*(*D*|*θ*) and the statistics of observed data *S*(*D*\*); and (iv) accept *θ* if the distance *d*(*S*(*D*|*θ*), *S*(*D*\*)) is less than some threshold value *ε*. This process is repeated multiple times to obtain the approximated posterior distribution. ABC-SMC algorithm^1^ speeds up the standard rejection ABC by constructing a set of intermediate distributions, which are defined by a sequence of threshold values *ε*_*t*_ in decreasing order, *ε*_1_ > *ε*_2_ > … > *ε*_*T*_ ≥ 0. Each intermediate distribution is generated from the previous distribution using a sequential importance sampling scheme. The number of accepted particles and thresholds *ε*_*t*_ must be provided by the user.

### ABC with Gaussian process (GP) emulation

To reduce the computational cost of running a large number of simulations in ABC, a GP emulator is first constructed to quantify the mapping from model parameter *θ* to the aforementioned distance *d*:

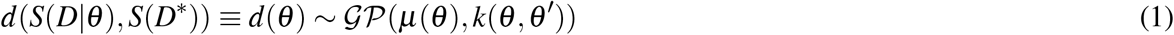

This GP is (re-)trained based on simulations of a relatively small number of parameters *θ* = [*θ*_1_, …, *θ_n_*]^*T*^, referred to as design points. The design points are sampled from the prior, or the posterior of the previous step in ABC-SMC, and selected in a way to control both the emulation accuracy and computational efficiency. Prediction of the distance for other particles can then be obtained from the GP posterior, without simulating the model. The GP emulation and re-training process can then be used with either rejection ABC or ABC-SMC algorithms, as shown in Figure 1.

**Figure 1.**
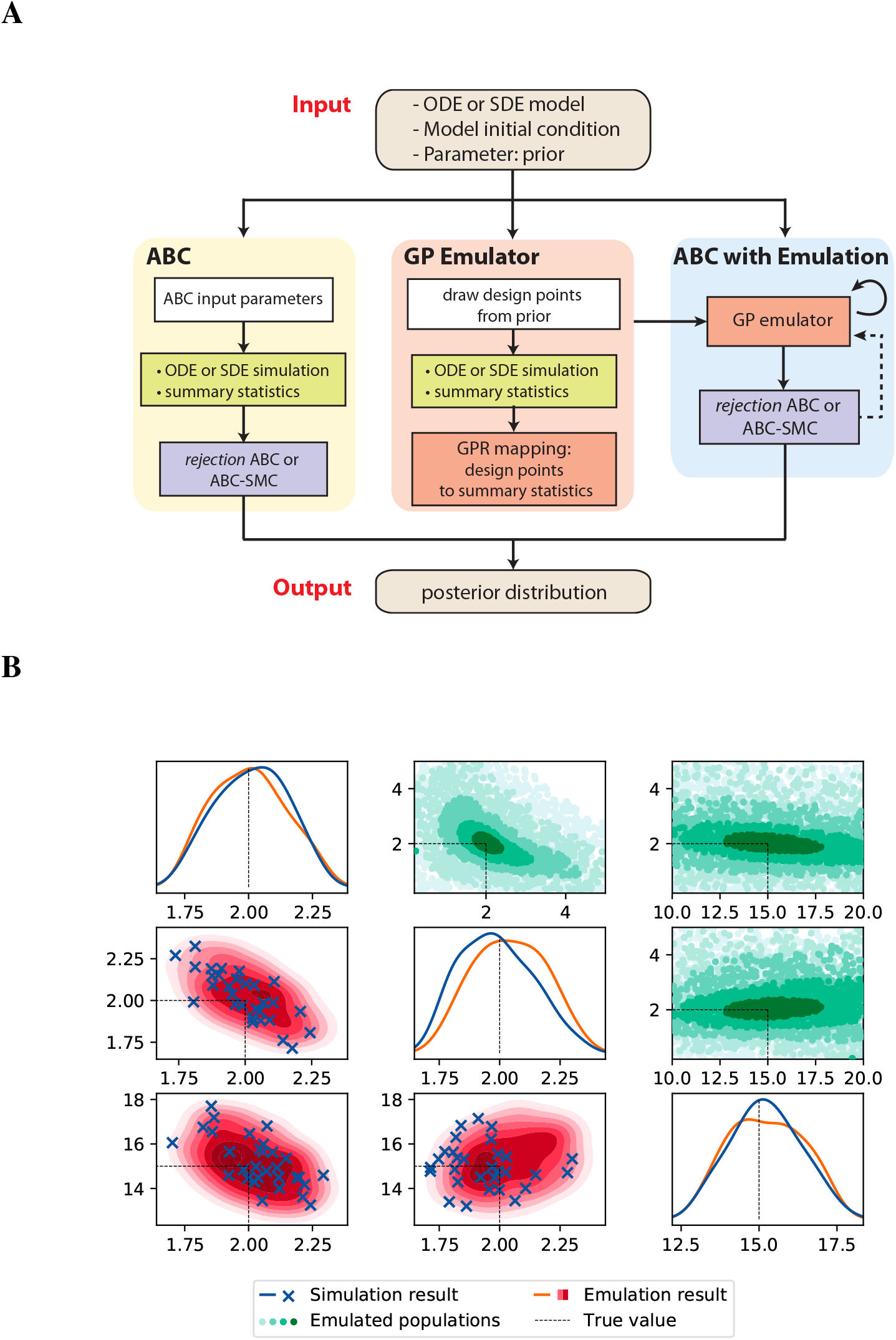
(**A**). Schematic diagram of GpABC package structure. The software can perform either purely Monte-Carlo simulation based ABC (i.e. rejection ABC, or ABC-SMC), or computational efficient ABC with emulation, where the GP emulator is first (re-)trained based on simulation from selected design points in the prior. Dashed arrow indicates the emulator re-training can be part of the ABC-SMC algorithm as design points are selected from different SMC populations iteratively. (**B**). Parameter inference results of the deterministic three-gene model^9^ using ABC-SMC algorithm. Subplots on the diagonal and lower triangular show marginal and joint posterior distributions of parameter estimates in the final ABC-SMC population (simulation in blue and emulation in red). Scatterplots above the diagonal show intermediate ABC-SMC populations with GP emulations; darker colour indicates decreasing threshold.

### Stochastic simulation and model selection

Biochemical reactions are stochastic in nature, and the distribution of stochastic simulation trajectories is generally non-Gaussian. To meet the Gaussian noise assumption of a GP and to consider computational efficiency, we employ the linear noise approximation (LNA), a first-order expansion of the stochastic differential equation. Users can select whether to perform ABC or ABC emulation for deterministic or stochastic modeling. In addition to parameter inference, a model selection algorithm^1^ is also implemented, where model indicators *m* are treated as an extra parameter. The joint posterior distribution over the combined space of models and parameters *p*(*m*, *θ*|*D*) can be obtained via a ABC-SMC scheme similar to the parameter inference, and finally the *p*(*m*|*D*) is obtained by marginalizing over parameters.

### Package overview and features

Users can easily choose or define several parameters of the algorithm. For ABC, these are the summary statistics *S*(*D*|*θ*) or a subset of the model’s outputs, prior distributions, distance function *d*, the number of accepted parameters in the posterior, and threshold values *ε*_*t*_. The latter strongly depend on the dynamics of the biochemical process and the noise level. Users can also choose how to select the design points in the emulator re-training process (with several optional strategies), which is a trade-off between the emulator accuracy and the computational cost. GpABC provides users with information about the progress of ABC: accepted number of samples from each Monte-Carlo perturbation.

Package outputs include posterior populations of accepted parameters for each threshold value, as well as distances, *d*, for each accepted parameter. Whenever emulation is used, additional information about the GP emulator is also provided. The package provides plots of marginal and joint distributions of accepted parameters.

Performance depends on how well the model fits the data and the choice of thresholds *ε*. GpABC will keep exploring the parameter space until the necessary number of parameters is accepted, or the maximum number of attempts is reached. In simulation mode, the model is simulated on each attempt, so the cost of simulating the model has crucial impact on performance. In emulation mode, the model is simulated only for training the emulator; subsequently model emulation is done in batches. Training the GP has computational complexity 𝒪(*n*^3^), and emulating the model has complexity of 𝒪(*bn*), assuming batch size *b*.

## Summary

GpABC is a user-friendly and extendable Julia package that can perform standard simulation based ABC or ABC with GP emulation, for both deterministic or stochastic systems biology models. The package can be used to infer the posterior parameter distribution or select the best model structure that represents the data from candidate models.

## Acknowledgements

We thank the members of the Theoretical Systems Biology Group at Imperial College London for helpful discussions and enthusiastic support.

